# C-terminal lysine residues localise NLRP10 at lipid compartments and govern NLRP10 oligomer formation

**DOI:** 10.1101/2025.11.27.690698

**Authors:** Timo-Daniel Voss, Christoph Winterberg, Adrian Beck, Clarissa Gottschild, Leonie Mueller, Selina M. Enayat, Matthias Geyer, Thomas A. Kufer

**Author notes:** Corresponding author: Timo-Daniel Voss, PhD, Department of Immunology, Institute of Nutritional Medicine, University of Hohenheim, 70599 Stuttgart, Germany.

## Abstract

NLRP10 is an atypical member of the NOD-like receptor (NLR) family because it lacks a leucine-rich repeat domain at its C-terminus. Recently, NLRP10 has been shown to form an inflammasome, however, the mechanism and the trigger of activation remain elusive. Here, we show that in human epithelial cells and keratinocytes NLRP10 oligomerises in response to the stressor *m*-3M3FBS. NLRP10 co-localises with ASC upon overexpression, but ASC nucleation and recruitment were different to other NLRPs. While neither ATP hydrolysis nor the pyrin domain were required, the C-terminal tail region (aa 584-655) was both necessary and sufficient for oligomerisation. Generation of chimeric proteins showed that the function of the tail is conserved between human and mouse NLRP10 in respect to aggregation but convers different protein stability. Changes in the subcellular localisation of NLRP10 and oligomerization were dependent on the presence of evolutionarily conserved lysine residues in the tail region, which localise the majority of NLRP10 at lipid droplets. Our study identifies the C-terminal basic tail of NLRP10 as a key regulatory element for oligomerisation and localisation at lipid interfaces. These findings underline differences of NLRP10 activation in respect to other inflammasome forming NLRPs and suggest a role of lipids in NLRP10 activation.

## Introduction

The family of NOD-like receptors containing a pyrin domain (NLRPs) consists of multiple proteins in mammals. NLRPs are characterised by a tripartite domain architecture composed of an N-terminal effector pyrin domain (PYD), a central regulatory NACHT domain (a nucleotide-binding and oligomerisation domain present in NAIP, CIITA, HET-E, and TEP1), and usually a C-terminal leucin rich repeat (LRR) sensory domain. NLRPs of the immune system act as intracellular pattern recognition receptors (PRRs) and detect various damage-and pathogen-associated molecular patterns (DAMPS/PAMPS).

Among the NLRPs, NLRP10 is a unique member as it lacks a LRR domain and shows a non-myeloid expression pattern^1–3^. Together with NLRP1^4^, the highest expression of NLRP10 is found in human and mouse skin, especially in keratinocytes^3,5^. Genome wide association studies linked NLRP10 to atopic dermatitis^6^ and NLRP10 was shown to contribute to contact hypersensitivity^5,7^. In addition to this, NLRP10 mRNA and protein levels are upregulated in psoriasis^8^ and downregulated in atopic dermatitis^9^. Together, these findings suggest that NLRP10 plays a role in maintaining skin barrier homeostasis. Whether these roles of NLRP10 are mediated by a pattern recognition function is controversially discussed in view of the absent LRR domain, which is usually involved in DAMP/PAMP recognition. First reports suggested a role of NLRP10 as negative regulator of the NLRP3 inflammasome^1,2^ where the NACHT domain supresses ASC activation and subsequent IL-1β release^1^. In contrast, it was found that NLRP10 can also enhance NF-κB-mediated inflammatory responses towards the bacterial pathogen *Shigella flexneri* by binding and destabilization of the anti-inflammatory regulator Abin-1^3,10^. More recently, it has been shown that both mouse Nlrp10 and human NLRP10 can form inflammasomes and induce IL-1β release in epithelial cells^11,12^. These two studies showed that NLRP10 is activated by 2,4,6-trimethyl-N-(*m*-3-trifluormethylphenyl)benzene-sulfonamid (*m*-3M3FBS), an inducer of mitochondrial stress. Notably, a protein sequence alignment of human and mouse NLRP10 proteins highlights differences in their amino acid composition, with 69% sequence similarity^13^. The structures of the human and mouse NLRP3 PYD domains were determined by NMR spectroscopy and docking analyses indicated that the two domains may use different surfaces of the PYD to interact with ASC^14^. This challenges the paradigm that the two proteins have the same function; a feature similarly observed, e.g., for NLRP1^15,16^.

In the present study, we characterised the role of both human and mouse NLRP10 in response to *m*-3M3FBS-induced cellular stress. We found that evolutionarily conserved positively charged amino acids in the C-terminal basic tail region, direct the association of NLRP10 at lipid droplets and governs the formation of higher-order oligomers. Our study helps further understanding the role of NLRP10 and paves the ways for the search of natural activators thereof.

## Results

### Oligomerisation of NLRP10 upon *m*-3M3FBS treatment is ASC- and ATP hydrolysis-independent

In the first report about an NLRP10 inflammasome, the mitochondrial damaging compound *m*-3M3FBS was used to induce NLRP10 inflammasome formation^12^. However, the mechanistic details of NLRP10 activation remained elusive. To further explore how NLRP10 is activated and forms inflammasomes, we generated stable HeLa cells that allow expression of N-terminally eGFP-tagged human NLRP10 under the control of a tetracycline-repressor (Suppl. Fig. S1A). We used HeLa cells as they are a well-established cell biological model for studying the subcellular localisation of NLR proteins^17,18^. To monitor *m*-3M3FBS-induced mitochondrial damage in live imaging, cells were co-transfected with mScarlet fused to the mitochondrial pre-sequence of human cytochrome c oxidase subunit VIII^19^. We observed that *m*-3M3FBS induced damage of mitochondria within minutes marked by the release of mScarlet into the cytosol (Fig. 1A). Subsequently, NLRP10 aggregated in the cytoplasm ∼4 min after *m*-3M3FBS was applied. Thereby NLRP10 formed cytosolically dispersed circular structures rather than single specks, typically seen for other inflammasomes such as NLRP3^20^. We did not observe a distinct co-localisation of NLRP10 with mScarlet on the outer mitochondrial membrane marker TOMM20 (Suppl. Fig. S1B). To exclude that the absent formation of NLRP10 specks was related to low expression of ASC in HeLa cells^21^, we generated stable HeLa cells expressing ASC-RFP^22^ controlled by a mutated cytomegalovirus promoter (CMVd1), and transiently co-transfected these cells with eGFP-NLRP10. Upon *m*-3M3FBS treatment the cells showed the formation of both, NLRP10 aggregates and ASC specks, but only some NLRP10 aggregates recruited ASC (Fig. 1B). To support our findings, we treated the human keratinocyte cell line HaCaT with *m*-3M3FBS and performed indirect immunofluorescence using our NLRP10 specific antibody^3^. As for overexpressed NLRP10 we observed aggregation also for endogenous NLRP10 in HaCaT cells (Suppl. Fig. S1C) but failed to detect IL-18 release (Suppl. Fig. S1D). Next, to analyse whether ATP-binding of NLRP10 is important for the oligomerisation, we used the Walker A mutant K197A, which is conserved in the human NLRPs (Suppl. Fig. S2E,F) and abolishes NOD1-mediated functions of NLRP10^3^. Stable cell lines expressing this NLRP10 variant showed no difference in the NLRP10 aggregation dynamics upon *m*-3M3FBS treatment compared to wild type NLRP10 (Fig. 1C). Collectively, our data suggest that NLRP10 aggregates independent of ATP hydrolysis function and does not form classical inflammasomes in our cell models.

**Fig. 1.**
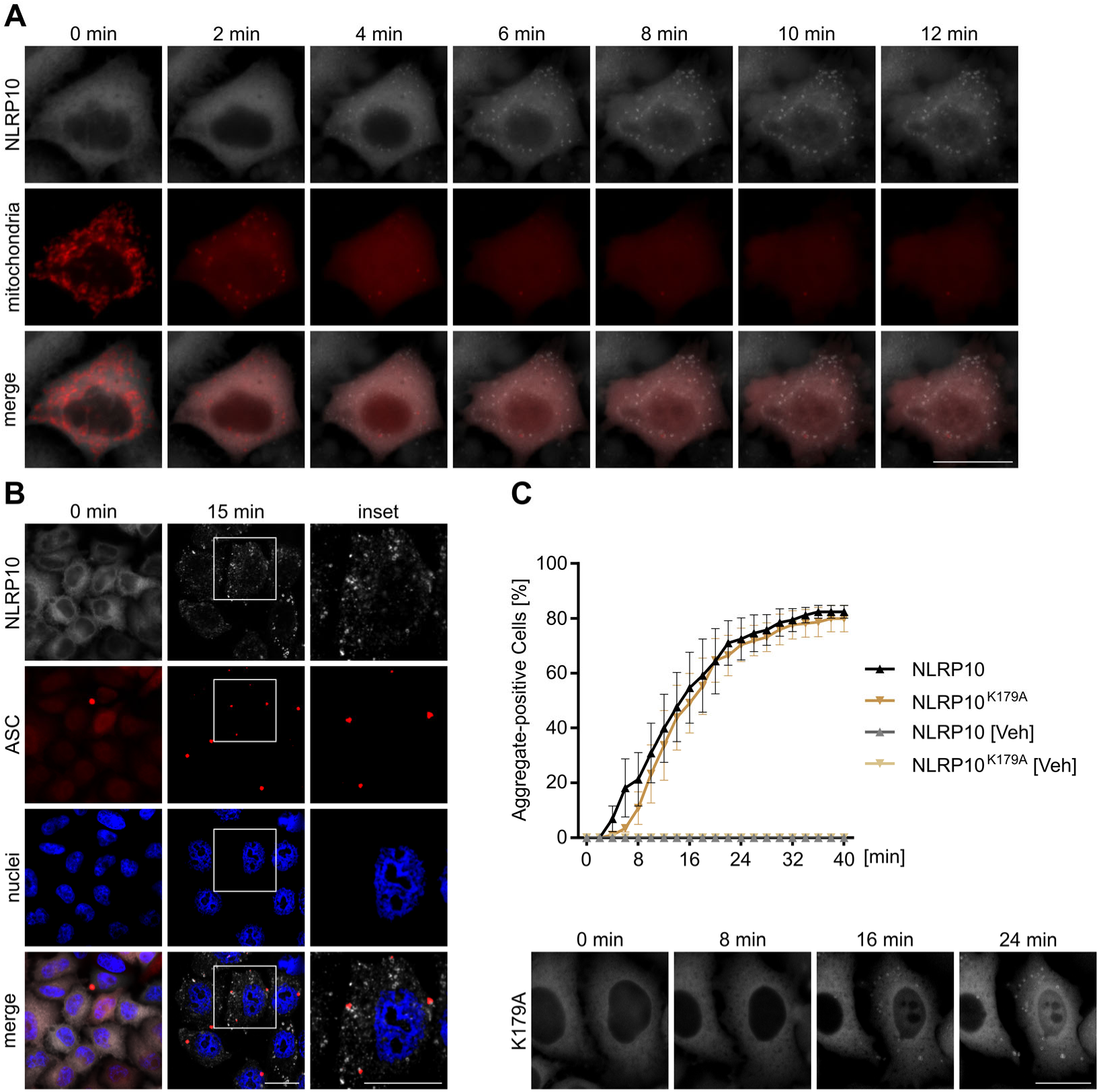
ASC and ATP-binding independent oligomerisation of NLRP10. (**A**) Live-cell imaging micrographs of eGFP-NLRP10 (white) and mts-mCherry (red) expressing HeLa cells treated with *m*-3M3FBS. (**B**) Micrographs of HeLa cells expressing eGFP-NLRP10 (white) and ASC-RFP (red) treated with 85 µM *m*-3M3FBS, nuclei were stained with HOECHST dye. (**C**) Quantification of the aggregation kinetics of NLRP10 or NLRP10^K179A^ with representative micrographs below. [Veh] referes to cells treated with *m*-3M3FBS solvent only. Scale bar: 20 µm. N=6 (C).

### NLRP10 aggregation depends on its C-terminal tail region

To further understand the aggregation process of NLRP10, we next investigated the role of various domains of NLRP10 considering evolutionary conservation. Super-positioning of protein structure predictions of NLRP10 from *Homo sapiens* and *Mus musculus* showed a high degree of structural similarity (Suppl. Fig. S2A). In contrast to known inflammasome-forming NLRPs such as NLRP3 or NLRP6, NLRP10 lacks a linker region between the PYD and the NACHT domain, which is encoded by an additional exon in NLRP3^14,23^, most likely resulting in a conformationally restricted flexibility of the NLRP10 PYD (Suppl. Fig. S2B).

NLRP10 lacks the typical LRR at the C-terminus; however, both human and mouse NLRP10 contain a protein region following the conserved helical domain 2 (HD2) of the NACHT (hereafter referred to as “tail”). This tail region is of low sequence complexity and no plausible structure prediction could be obtained for neither the human nor the mouse NLRP10 sequence classifying it as an intrinsically disordered region (IDR). To understand how the C-terminal tail and the PYD may influence NLRP10 complex formation we generated deletion constructs of both human and mouse NLRP10 lacking either of these domains (Fig. 2A). Stable HeLa cells expressing either of these NLRP10 truncation proteins, N-terminally fused to eGFP, were analysed for their propensity to form aggregates upon *m*-3M3FBS treatment in live-cell imaging (LCI). Deletion of the PYD did neither change the appearance of aggregates in the cell (Fig. 2B) nor their aggregation kinetics (Fig. 2C; black vs lavender). By contrast, deletion of the tail region (aa 584-655) or of the HD2–tail subdomains (aa 584-655) completely inhibited formation of NLRP10 aggregates (Fig. 2B,C). Next, to analyse the role of the tail region in murine Nlrp10 we expressed eGFP-tagged murine Nlrp10 as well as deletion constructs lacking the PYD or the C-terminal tail in HeLa cells lacking endogenous NLRP10 expression (Fig. 2D,E, Suppl. Fig. S2C,D). Murine Nlrp10 aggregated in response to *m*-3M3FBS treatment with comparable kinetics observed for human NLRP10 (Fig. 2D,E; black). As seen for human NLRP10, deletion of the tail region almost completely inhibited the formation of Nlrp10 aggregates, while deletion of the PYD only did not affect aggregation (Fig. 2D,E; lavender, magenta).

**Fig. 2.**
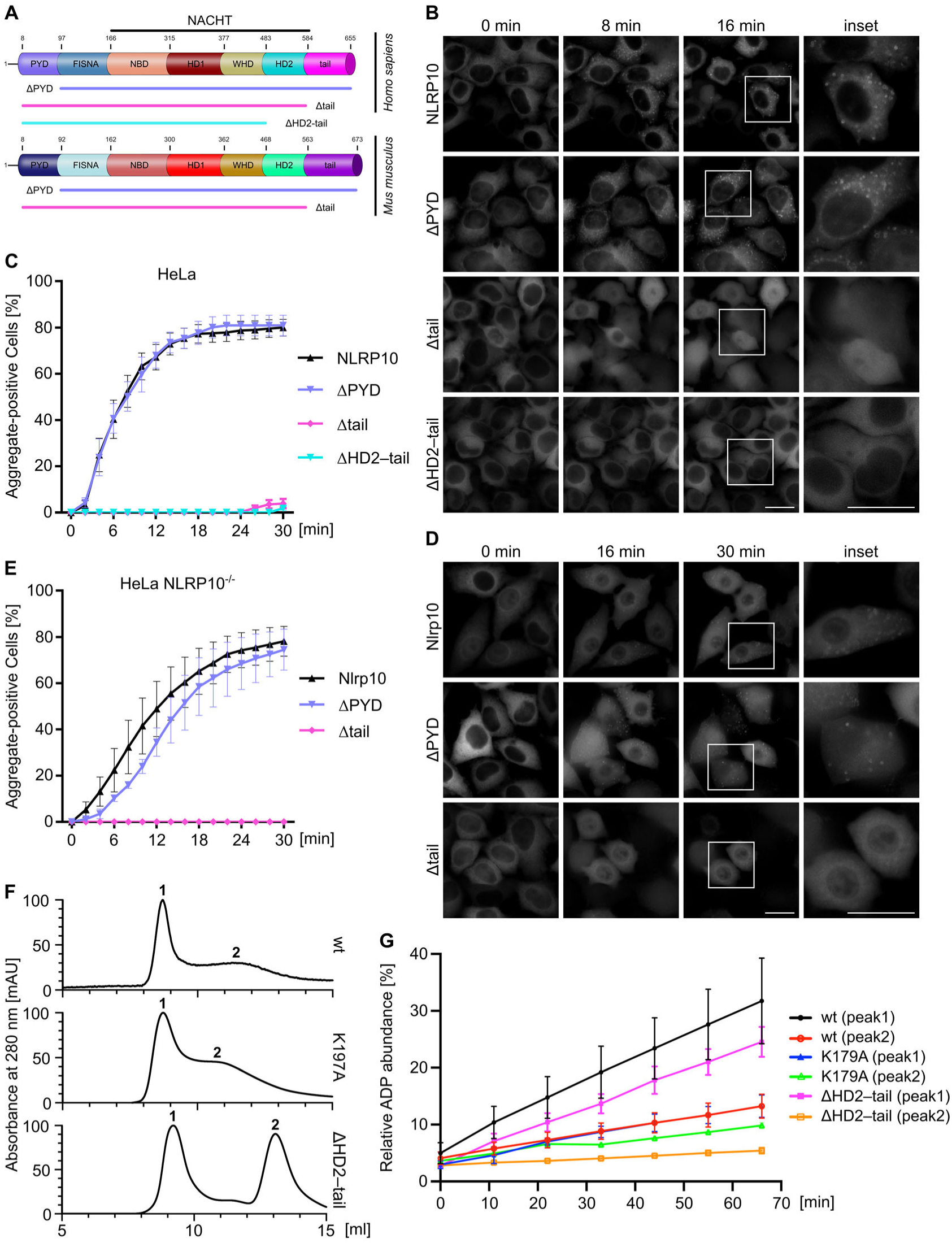
NLRP10 oligmerisation depends on C-terminal tail in human and murine NLRP10. (**A**) Schematic representation of Homo sapiens and Mus musculus NLRP10 and truncation mutants thereof. (**B, D**) Live-cell imaging micrographs of stable HeLa (**C**) or NLRP10-knockout HeLa (**E**) cells treated with m-3M3FBS (85 µM) expressing human (**C**) or murine (**E**) NLRP10. (**C, E**) Quantification of the aggregate-positive cells from (**B**) and (**D**). (**F**) Size exclusion chromatography absorbance spectrum of NLRP10 (4-655), K197A mutant or NLRP10ΔHD2–tail (aa 4-482) at 280 nm. (**G**) Relative ADP abundance over time as a measure of the ATP hydrolysis rate of peaks 1 and 2 measured by ion-paired reverse phase HPLC from (**F**). Scale bar: 20 µm. N=7-12 (**B**) or N=4-8 (**D**).

Seeing that the tail segment was required for oligomerisation, we further explored, if ATP-binding, which is required for an active protein conformation, is also compromised by the deletion of the tail. A construct with a mutated Walker A site (K179A) was used as control. The corresponding recombinant NLRP10 proteins were expressed in *Sf9* insect cells and purified to homogeneity. Proteins were separated by size exclusion chromatography and the two most abundant fractions, peak 1 (void) and peak 2 (NLRP10 oligomers), were collected (Fig. 2F). Subsequently, ATP hydrolysis was determined by measuring ADP production via ion-paired reverse phase HPLC analysis. The peak 1 fraction of wild type NLRP10 showed ATP hydrolysis activity, whereas the peak 2 fraction showed an about 3-fold reduced hydrolysis rate (Fig. 2G, black and red lines), in line with the assumption of an autoinhibited conformation as similarly observed for NLRP3^24^. As expected, the NLRP10 K179A mutant exhibited a significantly reduced ATP turnover rate, both in the peak 1 and peak 2 fractions (Fig. 2G, blue and green lines). The ΔHD2–tail variant again showed ATP hydrolysis activity in peak 1 almost to the same extent as the full-length NLRP10 protein (Fig. 2G, magenta line). Unexpectedly, deletion of the HD2–tail region resulted in NLRP10 monomer formation and almost completely blocked ATP hydrolysis (Fig. 2F and G, orange line). Collectively these data show that both human and murine NLRP10 can form dispersed cytosolic aggregates, which depend on the C-terminal tail region while the PYD and ATPase functionality are not required for this feature. This assigns a novel regulatory function to the tail region of NLRP10 that is conserved between mice and men.

### Evolutionarily conserved basic residues in the C-terminus of NLRP10 mediate oligomerisation

The observation that the tail regions of both human and murine NLRP10 are necessary for oligomerisation suggests an evolutionarily conserved function of this segment. However, the tail sequence in mouse Nlrp10 is longer compared to its human counterpart (112 aa vs 72 aa). Overall, NLRP10 seems to have underwent complex evolution in primates (Suppl. Fig. S3A), while Myomorpha Nlrp10 evolution is largely represented by the phylogenetic variance (Suppl. Fig. S3B). Among primates, the human NLRP10 homologue appears to have one of the smallest tail regions, only Squirrel monkey (*Saimiri boliviensis*) NLRP10 has a shorter tail (Suppl. Fig. S3A and Suppl. Table S1). Within the Cattharines, the NLRP10 sequence of Cercopithecidae separate from a mixed cluster of NLRP10 sequence homologues from Hylobatidae and Hominidae, while the human NLRP10 early diverged from this cluster (Suppl. Fig. S3C).

Sequence analysis showed that the human tail has a higher isoelectric point (pI ∼9.5) compared to mouse (pI ∼8.8) (Suppl. Table S1, Suppl. Table S2). However, the part of the murine tail that aligns with the human sequence (aa 566–622) has a very high pI of ∼10.1, whereas the remainder of the C-terminal sequence has a pI of ∼4.7 (Fig. 3K). The basic character of the tail sequence in human NLRP10 results from twelve lysine residues and one arginine, loosely distributed over the entire length of the sequence, which are opposed by only seven glutamates and one aspartate. This positive net value establishes the intrinsically disordered C-terminal region of NLRP10 as ’basic tail’, which is unique within the family of NLRPs.

**Fig. 3.**
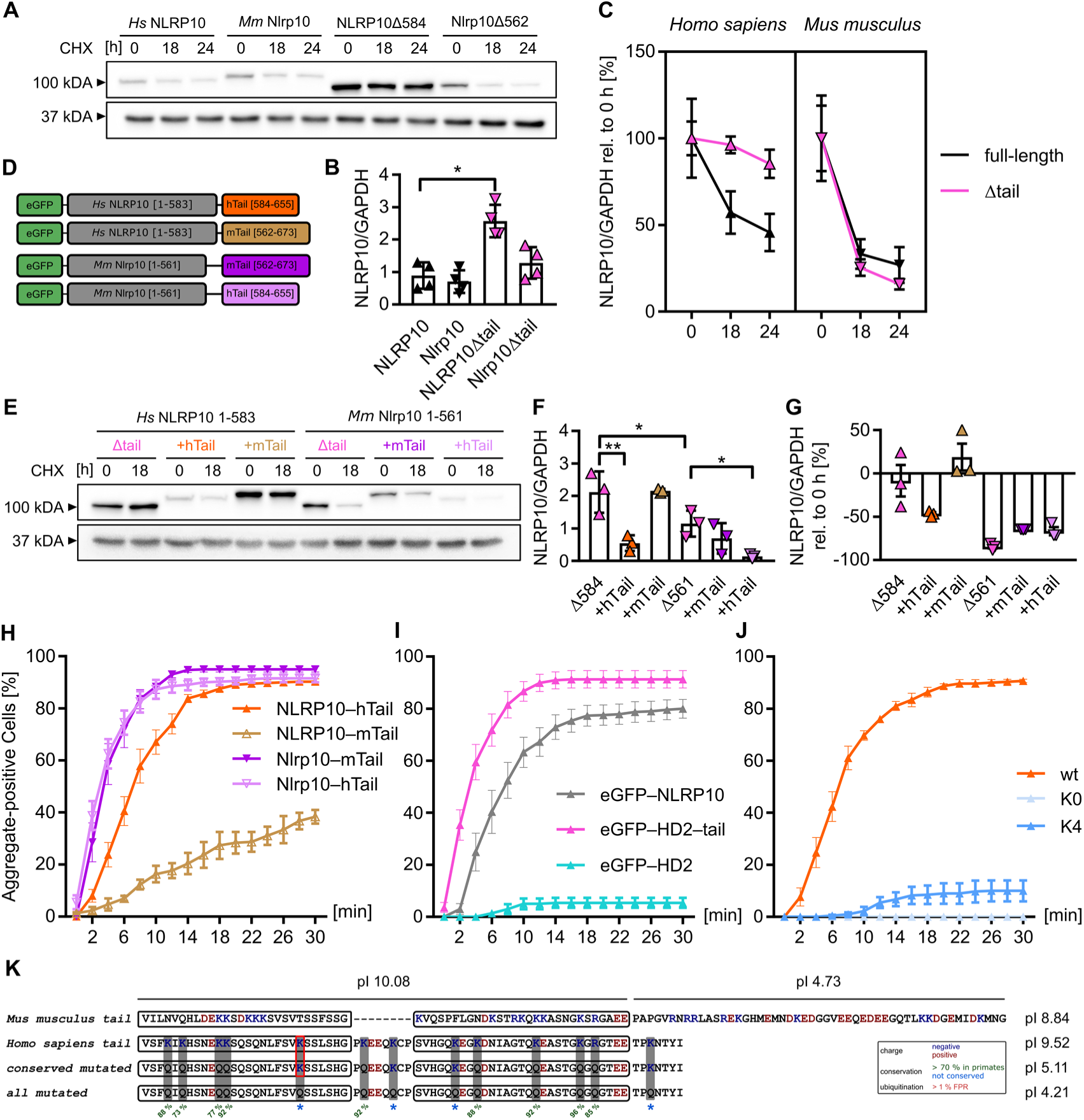
The basic tail of human NLRP10 governs protein stability and oligomerisation, which is dependent on lysine residues. (**A, E**) NLRP10 immunoblot image of transiently transfected HeLa cells to analyse the stability of various truncation mutants and chimeric NLRP10 proteins. Cells were treated with CHX (20 µg/ml) 48 h post-transfection (hpt) and lysed after 18 h or 24 h of CHX treatment. (**B, F**) NLRP10 levels at baseline (48 hpt) normalised to GAPDH as reference. (**C, G**) relative change of the protein levels in response to 18 h (**C, G**) or 24 h (**C**) CHX treatment. (**D**) Schematic representation of the chimeric NLRP10 variants used in (**E-H**). (**H-J**) Quantification of the aggregate-positive cells in live-cell imaging micrographs of HeLa cells treated with *m*-3M3FBS (85 µM) transiently expressing chimeric NLRP10 variants (**H**), NLRP10 tail (**I**) or NLRP10 lysine substituted variants (**J**). (**K**) Alignment of the *Mus musculus* and *Homo sapiens* NLRP10 C-terminus and mutated C-terminus of the latter. Positive (red) and negative (blue) amino acids are marked. Putative ubiquitination sites are represented by red boxes, conservation was determined with Jalview using all primate NLRP10 sequences. N=3 (**H**).

N- and C-terminal regions are often involved in the regulation of protein stability^25,26^. Prediction of ubiquitination consensus sites (Rapid UBIquitination prediction) revealed that the human tail has one putative high-confidence ubiquitination site (K605), while no high-confidence site was found in the mouse Nlrp10 tail. We expected that the proteins lacking the C-terminus should be more stable than the full-length proteins. Indeed, both human and mouse NLRP10 lacking the basic tail region (NLRP10Δ584 and Nlrp10Δ562) showed higher protein expression levels in HeLa cells after transfection with equal amounts of expression plasmid. Deletion of the human tail led to a 2.9 ± 0.6-fold increase in protein stability and deletion of the murine tail resulted in a 1.8 ± 0.7-fold increase, respectively (Fig. 3A,B). When blocking protein *de novo* synthesis by cycloheximide upon 18 h, NLRP10 was degraded to 57.1 ± 12.2 % while deletion of the tail led to a reduction of only ∼4 %. Murine Nlrp10 also showed a high turnover and protein levels decreased to 33.2 ± 8.5 % within 18 h. However, the deletion of the tail in mouse Nlrp10 did not affect its degradation (Fig. 3C).

Next, to corroborate the role of the basic tail from murine and human NLRP10 in protein stability, we generated chimeric constructs consisting of the human and murine core (PYD and NACHT) and the murine or human tail regions, respectively (Fig. 3D). When using the human core fused to the murine tail, we observed increased protein stability nearly to levels obtained for the tail deletion mutant. By contrast, with its intrinsic human tail the protein was less stable (Fig. 3E-G). Vice versa, when using the murine core, fusion to the human tail drastically destabilised the protein (∼20 % compared to murine tail). This strongly suggest that the human tail region contributes to protein turnover.

To understand if the human and murine NLRP10 basic tail regions are compatible with the core protein for oligomer formation, we transiently transfected HeLa cells with the four chimeric NLRP10 variants and measured NLRP10 oligomer-positive cells upon *m*-3M3FBS exposure as described above. The constructs containing the mouse PYD–NACHT domains showed aggregation at a comparable kinetics, regardless of the origin of the tail region (Fig. 3H and Suppl. Fig. S4A). The human NLRP10 core with the human tail aggregated with a similar kinetics as the murine proteins, as expected from our results using stable human and mouse NLRP10 expression (Fig. 2C,E). Unexpectedly fusion of the murine tail to the human core protein drastically reduced aggregation, only leading to 38.3 ± 4.5 % at 30 min (Fig. 3H), even within 60 min only half of the cells showed aggregates (53.7 ± 5.0 %) (Suppl. Fig. S4B).

Collectively, the highly conserved C-terminal tail of NLRP10 is necessary for protein aggregation and stability. To test if the basic tail region is sufficient for aggregation we generated eGFP proteins with N-terminal fusion to the HD2 domain of human NLRP10 with and without the tail region and transiently transfected HeLa cells with these constructs. Cells were subsequently treated with *m*-3M3FBS and the aggregation behaviour monitored. Cells expressing the eGFP-HD2–tail construct immediately responded with aggregate formation (Fig. 3I), which resembled the aggregates of NLRP10 (Suppl. Fig. S4C). Cells that only expressed the HD2 domain did not respond with aggregate formation in response to *m*-3M3FBS, confirming that the tail is necessary and the HD2–tail sufficient for *m*-3M3FBS-induced oligomerisation. The unusually high pI of the tail led us to hypothesise that the presence of conserved positively charged amino acids mitigates aggregate formation. To test this, we used human NLRP10 and mutated the arginine residue and either all twelve or only the nine highly conserved lysine residues (Fig. 3K) and evaluated the propensity of these proteins to aggregate upon *m*-3M3FBS treatment. Mutation of all lysine and arginine residues to glutamine (K0) completely inhibited the oligomerisation of NLRP10 in HeLa cells. Deletion of the highly conserved lysine residues only (K4) was sufficient to vastly inhibit aggregation to 10.0 ± 8.0 % of the cells compared to ∼80 % observed for NLRP10 (Fig. 3J), suggesting that these amino acids control oligomerization.

### The NLRP10 oligomers are located at lipid compartments

We found that upon *m*-3M3FBS treatment NLRP10 forms ring like structures and that basic residues in the tail region are required for oligomerisation. This led us to hypothesise that NLRP10 might be recruited by or to lipid compartments to form oligomers at the membrane bilayer. Consistent with this, we found lipid membranes co-purifying with NLRP10 protein expressed in *Sf9* cells, as shown by electron microscopy (Fig. 4A). In contrast, when purifying the NLRP10 K0 protein lipid membranes were only found to a very low extent. To analyse co-localisation of NLRP10 with membranous cellular compartments, we used live cell imaging with mCherry expression constructs labelling cellular compartments including early endosomes, lysosomes or the endoplasmic reticulum (Suppl. Fig. S5). We observed that NLRP10 at best only partially co-localised with endosomal and lysosomal markers, consistent with the lack of a clear co-localisation of NLRP10 with the ER, endosomal, lysosomal or Golgi compartments in HEK293 cells^12^.

**Fig. 4.**
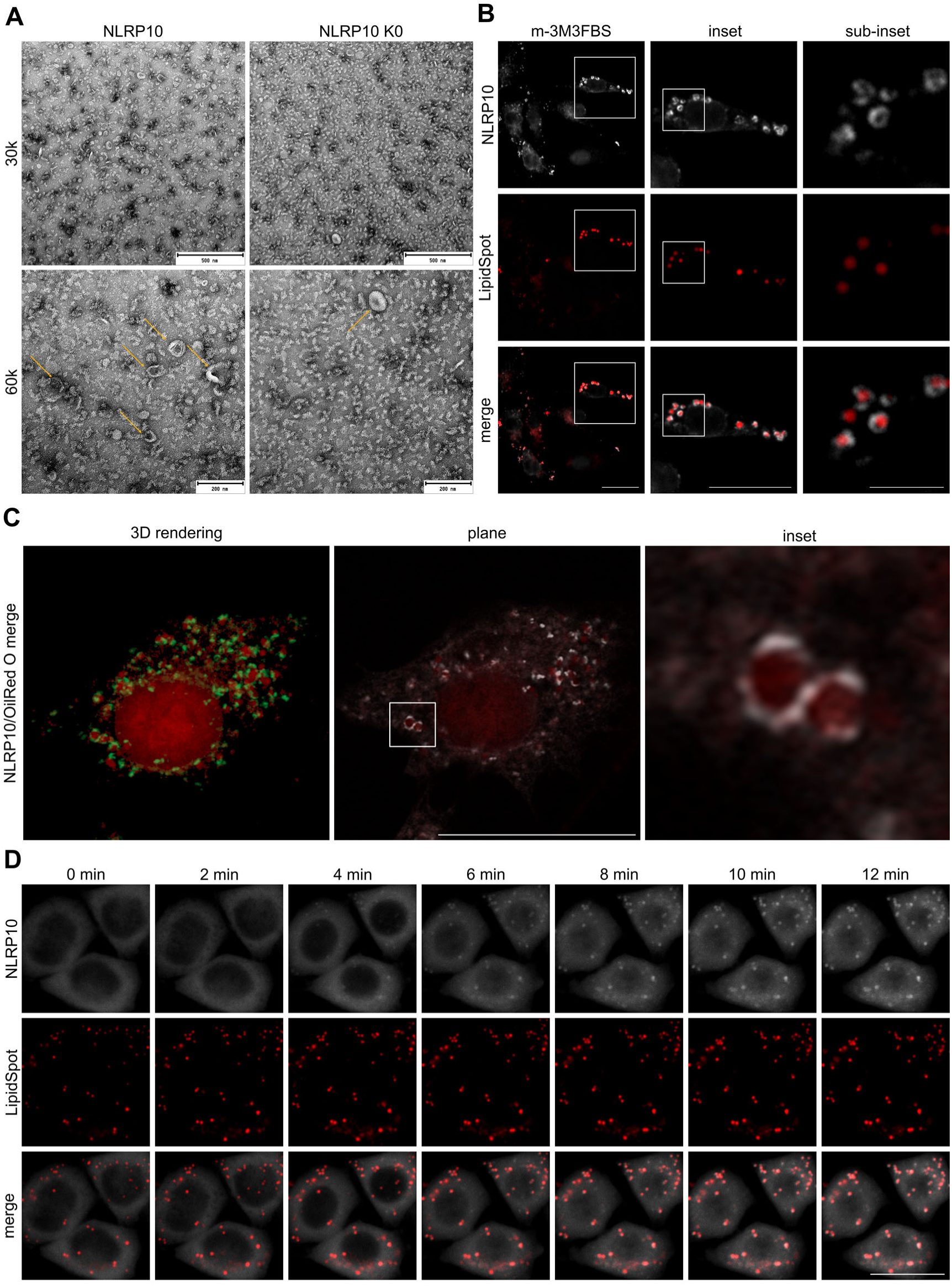
NLRP10 oligomers associate with membranes and lipid droplets. (**A**) Negative stain electron microscopy images of wild type NLRP10 or the NLRP10 K0 mutant (peak 1) with magnifications of 30k and 60k. Arrows indicates membranes at 60k. (**B**) Micrographs of HeLa cells expressing eGFP-NLRP10 (white), treated with *m*-3M3FBS (85 µM). Lipid droplets were stained with LipidSpot™ 610. (**C**) 3D rendering image of eGFP-NLRP10 (green/white) expressing HeLa cells stained with OilRedO (red) upon treatment with *m*-3M3FBS. (**D**) Live-cell imaging micrographs of eGFP-NLRP10 (white) expressing HeLa cells stained with LipidSpot™ 610 (red) upon treatment with *m*-3M3FBS. Scale bar: 500 nm (A, upper panel) and 288 nm (lower panel), 20 µm (B, left and middel panel, C) or 5 µm (B, right panel).

Inspired by the shape of the NLRP10 structure, we next tested if NLRP10 might be recruited to lipid droplets, cytosolic lipid storage compartments and immune hubs important for integrating energy metabolism and host defence^27,28^. To this end, we treated our stable eGFP-NLRP10 expressing cells with *m*-3M3FBS and stained fixated cells with OilRed O or LipidSpot™ to stain lipid droplets. Here, NLRP10 formed oligomers, which localised at lipid droplets (Fig. 4B). Confocal imaging and 3D rendering showed that NLRP10 clearly localised around lipid droplets (Fig. 4C). Live-cell imaging revealed that the lipid droplets are frequent in the cells and only upon *m*-3M3FBS treatment NLRP10 oligomer formation started at these structures (Fig. 4D). Taken together, our data showed that while we observed recruitment of NLRP10 to lipid droplets, identifying a novel role of lipid droplets in the activation process of NLRP10.

## Discussion

Human NLRP10 is implicated in inflammatory modulation, however in contrast to other inflammasome forming NLRPs such as NLRP3, it is virtually absent in myeloid cells^3^. At physiologically relevant levels it is expressed in keratinocytes^3^. This makes keratinocytes and epithelial cells ideal models for studying the function of NLRP10. It is likely that NLRP10 contributes to skin homeostasis. This is supported by a study showing that knockout of NLRP10 significantly reduces epidermal thickness in human skin equivalents^9^ as well as by our previous finding that NLRP10 expression in keratinocytes is associated with inflammation in type IV hypersensitivity in the skin^29^. Even though NLRP10 does not contain an LRR domain, its C-terminus consists of 72 amino acids distal of the NACHT domain. This segment, predicted to be intrinsically disordered, is a variation of the transition LRR (trLRR) domain in NLRs which encompasses for example in NLRP3 a 42 residue long acidic loop^30^. Inflammasome formation is not depended on the LRRs as shown for NLRP3, which can still be activated via nigericin to form the canonical inflammasome when the LRR is depleted^31^. It therefore came as no surprise that two recent reports suggested that NLRP10 can form an inflammasome^11,12^. While the first report was mostly focussing on expression in HEK293 and N/TERT cells, the second analysed the physiological role of Nlrp10 in DSS-induced colitis in mice and found a protective role of Nlrp10 expression^11,12^. If all these functions are associate with inflammasome formation of NLRP10 or might be mediated by inflammasome-independent functions of NLRP10, as we have shown in the context of bacterial infection^3^, awaits clarification. In our present study we focussed to work out NLRP10 intrinsic regulatory mechanisms for aggregate formation. In human HeLa cells and HaCaT keratinocytes we showed that NLRP10 forms oligomers upon activation and observed a cytosolic dispersed pattern of aggregated NLRP10, which is reminiscent with a dotted dispersed aggregation of NLRP10 seen in primary human keratinocytes^12^. However, we did not detect downstream IL-18 release. The function of the NLRP10 aggregates needs further clarification and these could be pre-forms of inflammasomes as observed for NLRP3^18^. Upon over-expression of ASC in our stable NLRP10 HeLa cells, we confirmed that NLRP10 can form stimulus-independent specks with ASC^12^ (Suppl. Fig. S6A). When we used stable HeLa cells expressing ASC-RFP controlled by a CMVd1 promoter, which has reduced expression levels compared to CMV^32^, ASC aggregated prior NLRP10 recruitment, indicating that NLRP10 is not the seed for ASC filament formation (Suppl. Fig. S6B). This, together with the lack of a flexible linker between the PYD and NACHT domains in NLRP10, strongly suggests that NLRP10 does not form a classical NLRP3-like inflammasome.

Our results demonstrate that the C-terminal tail is both required and sufficient for oligomerisation of NLRP10 (Fig. 2 and 3). This process is linked to changes in the sub-cellular localisation of NLRP10 aggregates within the cells. It was suggested that NLRP10 aggregates co-localise with mitochondria^12^. Using our eGFP-NLRP10 construct or immunostaining of endogenous NLRP10 (Suppl. Fig. S6C), we could not detect a clear mitochondrial co-localisation in our cells. By contrast, we observed localisation of NLRP10 complexes at lipid droplets (OilRed O or LipidSpot-positive structures) and partial co-localisation with early endosomes and lysosomes. Furthermore, we observed an enrichment of lipid membranes in fractions of purified NLRP10 protein expressed in *Sf9* cells, suggesting that NLRP10 can bind to membranes (Fig. 4). Based on our observation that lysine residues in the tail of NLRP10 are needed for oligomer formation and sub-cellular reorganisation, we hypothesise that the tail mediates localisation at lipids. Indeed, we could show that a construct containing the HD2 and tail domains was sufficient to localise at membranes. Notably, a prerequisite for this was treatment of the cells with *m*-3M3FBS. We assume it unlikely that the HD2–tail has propensity to detect any triggers and thus propose that changes in the composition of membrane lipids are the actual signal for NLRP10 clustering. Support for this comes from a study on the role of lysine clusters in peptides that bind to the membranes with a sigmoidal dependence^33^, which matches our aggregation kinetics. Furthermore, NLRP3 harbours a lysine-stretch, which is important for its trans-Golgi localisation^18^. Additionally, we showed before that NLRP10 can localise to the plasma membrane of HeLa cells upon bacterial infection and is found in flotillin-2 positive membrane fractions when co-expressed with NOD1^3^. Taken together, we provide evidence that the basic tail region of NLRP10 anchors the protein at membranes.

Asking if this mechanistic feature of NLRP10 is evolutionarily conserved, we showed that also murine Nlrp10 forms aggregates towards *m*-3M3FBS. Even though the tail region of murine Nlrp10 is substantially longer, the aggregation was engaged in a similar manner. The functions of the tail regions were confirmed by chimeric proteins consisting of murine tail and human core NLRP10 fusions, and vice versa. Unexpectedly, the murine tail was not fully compatible with the human core as this chimeric construct showed reduced aggregation kinetics. This observation might be explained by core–tail interactions. Interestingly, the human NACHT domain contains a sequence (FEEKLKKRGL, aa 258-267) just following the Walker B motif, which is absent in mouse Nlrp10. This polar region is supposed to form an exposed loop, whose overall basic charge could possibly attract the extended acidic tail region of mouse Nlrp10 (Fig. 3K). On the other hand, this might be a species-specific effect and further studies will show if different stimuli induce NLRP10 aggregation in mouse and men. In Muroidea the evolution of NLRP10 has not been as diverse as in primates. Both, the sequence conservation as well as the length and pI of the basic tails, largely resembles the evolutionary conservation of the subfamilies, where all Creticidae and Muridae form a distinct cluster and both separate from *Jaculus jaculus* (Suppl. Fig. S3B and D). We cannot explain how the presumed divergence that changes the picture in primates has happened and if this has an influence on the function. However, the high conservation of the lysine residues implies that the localisation of NLRP10 at lipid compartments is conserved. The interesting finding that the mouse tail fused to the human core leads to slowed aggregation allows us to speculate that the prolonged tail might stabilise the autoinhibited state of the protein and it will be of interest to analyse other primate NLRP10 candidates in their aggregation behaviour including *Saimiri boliviensis* (shortest protein and tail), *Aotus nancymaae* (longest protein) and, e.g., *Pan paniscus* (longest tail). This would also help to understand the physiological role of NLRP10, its aggregation and localisation.

Taken together, our work identifies the C-terminal basic tail of NLRP10 as an important regulatory element for NLRP10 activation. Furthermore, we provide evidence that this domain clusters NLRP10 at lipid rich compartments in the cells. Linking NLRP10 to a ’sensing’-function of lipids will help identify the activation cues which likely converge at affecting lipid composition of cellular membranes. Moreover, our data emphasises the significances of lipids in the activation processes of NLRs, as demonstrated by the interaction of NLRP3 with PI(4)P and by the NOD1/2 interaction with sphingosine-1 phosphate (S1P)^34^. This will also provide novel insights into the role of NLRP10 in skin disease, such as atopic dermatitis, which is linked to profound alterations in cellular lipids and lipid modifying enzymes^35^.

## Methods

### Molecular Cloning

#### PCR

50 ng template DNA were mixed with primers (Suppl. Table S3) and Phusion High-Fidelity PCR master mix (#F531, Thermo Fisher Scientific; Waltham, MA, USA) according to the manufacturers protocol. PCR products were analysed by gel electrophoresis. PCR products were purified using the PCR/Gel purification kit (#740609, Macherey-Nagel; Dueren, GER) according to the manufacturers protocol and DNA was eluated in nuclease-free water.

#### Agarose gel electrophoresis

To separate DNA fragments, 0.5 % agarose (Carl Roth) in TAE buffer [121 g Tris, 28.5 ml acetic acid, 9.3 g 0.5 M EDTA, 500 ml H_2_O] gels were used. ROTI®GelStain red (#0984, Carl Roth; Karlsruhe, GER) was added directly in the agarose gel and separation was performed at 10 V/cm for 25 - 60 min. DNA was visualized with UV light (365 nm) using the Fusion FX Camera System (Vilber Lourmat). Size of DNA fragments was estimated by Gene Ruler 100 bp plus and GeneRuler 1 kb DNA ladders (#SM0321 and #SM0311, TFS) as standards.

#### Restriction digest

PCR products and vectors were cloned using FastDigest restriction enzymes with 10x FastDigest (FD) green buffer (both TFS) in a total volume of 20-30 μl at recommended conditions.

#### DNA ligation

T4 DNA ligase (#EL0011, TFS) was used as recommended in a total volume of 20 μl for 10 min at room temperature followed by 5 min heat-inactivation at 70 °C. Digested and purified insert was applied in a 1:3 mass/size ratio to the digested and purified expression vector.

#### Mutagenesis

The K179A mutation of NLRP10 has been published previously^3^. Plasmids carrying the NLRP10 tails with deleted lysine and arginine residues were generated at Eurofins genomics and then sub-cloned into NLRP10Δ584 plasmid in a pcDNA5/FRT/TO backbone with deleted stop codon.

#### Transformation and DNA isolation

For amplification of mammalian expression plasmids, 10 μl of the ligation mix was combined with 50 μl chemically competent *E. coli* DH5α [F-Φ80lacZΔM15 Δ(lacZYAargF) U169 deoR recA1 endA1 bsdR17 (rk+, mk+) phoA supE44 thi-1 gyrA96 relA1 λ-]. Solution was gently mixed and incubated on ice for 30 min. Bacteria were heat-shocked for 30 sec at 42 °C and immediately placed on ice for 5 min before 1 ml pre-warmed LB was added to the bacteria. After incubation for 1 h at 37 °C and 400 rpm bacteria were pelleted at 3,000 x g for 5 min and 800 μl of the medium were removed. The bacteria were resuspended in the remaining 250 μl LB and spread on a LB agar plate containing 50 μg/ml kanamycin or 100 µg/ml ampicillin and incubated over-night at 37 °C. Single colonies were expanded in 7 ml LB containing antibiotics and DNA was extracted with the NucleoBond Xtra Midi Kit (#740410) or the NucleoSpin Plasmid Mini Kit (#740588; Macherey-Nagel), according to the manufacturer’s protocol. Plasmid DNA was eluted with endotoxin-free water. Plasmids were digested to test for successful integration of inserts and sequenced (Eurofins Genomics; Ebersberg, GER). Plasmids for organelle tracking [mCherry-Lysosomes-20 (kindly provided by Michael Davidson (RRID:Addgene_55073), mCherry-Sec61β kindly provided by Stephen Royle, (RRID:Addgene_172445)^36^, pFX-mCherry-EEA1 (kindly provided by Yusuke Ohba, (RRID:Addgene_174452)^37^, 4xmts-mScarlet-I (kindly provided by Dorus Gadella RRID:Addgene_98818)^19^].

### Generation of Stable HeLa Cell Lines

The maternal HeLa Flp-In cells were kindly provided by the Hentze Lab (EMBL Heidelberg) and allow doxycycline-inducible expression of various NLRP10 proteins sub-cloned into the pcDNA5/FRT/TO expression vector. To generate stable cells, 7.5 x 10^5^ maternal HeLa cells were seeded on a 6-well plate in 3 ml growth medium. The next day, medium was changed to 2.5 ml fresh growth medium and cells were co-transfected using Lipofectamine 2000 according to the manufacturers protocol. Briefly, 400 ng of the pcDNA5/FRT/TO (#V652020, TFS) with the respective insert were mixed with 3.6 µg pOG44 (#V600520, TFS) plasmid in 250 µl Opti-MEM™ I (#31985070, TFS). 10 µl Lipofectamine 2000 (#11668019, TFS) in 250 µl Opti-MEM were added to plasmid DNA and incubated for 20 min, mix was added dropwise to the cells. The following day, cells were detached and spread onto three 10 cm dishes in a total of 10 ml growth medium each. 72 h after spreading, selection antibiotics hygromycin (500 µg/ml) and blasticidin (10 µg/ml) were added and medium was changed every two-three days. Within ∼14 days single-cell colonies appear and they were transferred into 100 µl trypsin-EDTA in 96 well plates to allow cells to separate from each other. Cells were subsequently expanded in a 96 well, then 24 well and 6 well. Cells were then analysed for successful integration at the locus by fluorescence microscopy, immunoblot analyses and β-galactosidase assay. Single clone cultures were used for experiments.

### Live Cell Imaging (LCI) and Microscopy

HeLa Flp-In eGFP-NLRP10 cells were seeded on a glass bottom culture dish with four compartments (Greiner) at a density of 10^5^ cells per subdivision in 500 μl growth medium. Prior to *m*-3M3FBS (Tocris Bioscience, UK; MedChemExpress, NJ, USA) treatment and LCI, cells were washed with 500 μl FluoroBrite (Gibco, ThermoFischer Scientific) and then incubated in 400 μl FluoroBrite supplemented with 2 mM L-glutamine for 30 min at 37 °C, >95 % rH and 5 % CO_2_. Images were taken every 2 min over a time course of 30-60 min. LCI was performed at 37 °C using a Leica DMi8 microscope equipped with a climate chamber and an HC PL APO x63/1.40 oil objective. The Leica LAS X software was used to process the acquired images. Cells were regarded as aggregate positive at the time-point where multiple (>5) aggregates were visible, cells were counted manually by four different experimenters with at least 2 frames (30-60 cells per frame) per condition. The same microscope was used to image fixated cells.

For confocal imaging we used the Zeiss confocal laser scanning microscope LSM 900 with Airyscan 2 with Axio Observer 7. For 3D rendering deconvolution principles LSM Plus was used. The Zeiss microscope is a service by the Imaging Unit of the Core Facility Hohenheim (CFH-IMG) of the university of Hohenheim, Stuttgart.

### Protein Expression and Purification

The coding sequence of human NLRP10 (aa 4-655), Walker A mutant human NLRP10 (aa 4-655 K179A) or human NLRP10ΔHD2–tail (aa 4-482) was PCR-amplified and inserted into the pACEBac1 acceptor vector, carrying an N-terminal MBP-tev affinity purification tag. All constructs were expressed in *Sf9* insect cells using the baculovirus expression system. Cells were lysed in pre-chilled lysis buffer (wt 4-655 and mutant: 25 mM Tris pH 7.5, 300 mM NaCl, 5 mM ß-ME, 5% glycerol; 4-482: 20 mM Tris pH 7.8, 150 mM NaCl, 5 mM ß-ME, 0.5 mM ADP, 10 mM MgCl2) supplemented with DNase (30 µM) and PMSF (1 mM) and subsequently sonicated. The cleared lysate was subjected to MBP affinity chromatography using a MBPtrap column (GE Healthcare, Munich, Germany), equilibrated with lysis buffer. The protein was eluted in SEC buffer supplemented with 10 mM maltose. The protein was subsequently subjected to gel filtration chromatography using a Superose 6 Increase 10/300 GL or Superdex 200 Increase 10/300 GL column (GE Healthcare, Munich, Germany) in SEC buffer; wt 4-655 and mutant: 25 mM Tris pH 7.5, 300 mM NaCl, 5 mM ß-ME, 5% glycerol; 4-482: 20 mM HEPES pH 7.8, 150 mM NaCl, 1 mM TCEP, 0.5 mM ADP, 10 mM MgCl2, 150 mM l-arginine. Protein quality was analysed by SDS-PAGE and target containing fractions were pooled, concentrated and snap-frozen in liquid nitrogen for further analysis.

### Negative Stain EM

Negative stain electron microscopy was employed to evaluate protein sample quality, focusing on aggregation, heterogeneity and overall integrity. Carbon-coated copper grids were glow-discharged and then incubated for 1 min with 5 µl of the target protein sample. Excess sample was removed with blotting paper. The grid was washed by sequentially immersing it in three individual 20 µl drops of the corresponding purification buffer. Each time, residual liquid was removed. The grid was stained with 2 % uranyl acetate for 30 sec, after which the stain was carefully blotted-off and the grid was air-dried. Imaging was carried out on a JEOL JEM-2200FS TEM operating at 200 kV and equipped with a CMOS camera (TemCam-F416). Micrographs were collected at 30k and 60 k magnification.

### Ion-Paired Reverse Phase HPLC

Ion-paired reverse phase HPLC was used to analyse the amount of nucleotide released upon ATP hydrolysis reactions. Nucleotides were separated using a Chromolith Performance RP-18 endcapped 100–4.6 HPLC column and the corresponding guard cartridge Chromolith RP-18 (Merck, Darmstadt, Germany). The measurement was performed using an Agilent 1260 HPLC System (Agilent Technologies, Inc., Santa Clara, USA). The mobile phase was composed of: 10 mM tetrabutylammonium bromide, 30 mM K_2_HPO_4_, 70 mM KH_2_PO_4_, 0.2 mM sodium azide, 4% acetonitrile, pH 7.5. The analysed protein samples were diluted to a final concentration of 3 μM in SEC buffer and were incubated with 100 μM ATP (Jena Bioscience, Jena, Germany) at 25 °C. The samples were incubated in glass vials (Waters Corporation, Milford, USA). 10 μl aliquots were taken every 10 min with 1 min delay time. The HPLC measurement was performed at 25 °C at a flow rate of 1 ml/min and the absorbance was continuously detected at 280 and 254 nm, respectively. Additionally, a standard measurement for single nucleotides was performed. Peak integrals were determined with the according analysis software and normalized to the standard measurement. The shown hydrolysis data refers to n = 3 or n = 2.

### Immunoblot Analysis

Proteins were separated by SDS-PAGE and transferred to nitrocellulose membranes (Amersham™ Protran®). Proteins were detected by using mouse IgG1κ anti GFP antibody (Merck-Roche, #11814460001; RRID:AB_390913), mouse anti GAPDH (#sc-47724, Santa Cruz Biotechnology; RRID:AB_627678), HRP-conjugated goat anti mouse and rabbit IgG antibodies (#170-6516; RRID:AB_11125547 and #170-6515; RRID:AB_11125142) and detected them using the Clarity Western ECL Substrate (BioRad, #1705061). Signals were recorded on an electronic camera system (Vilbert Fusion FX).

### Densitometric Analysis

To analyse the densitometry of signals we used the open-source software Fiji (RRID:SCR_002285)^38^ and normalised to GAPDH signal as loading control and corrected by mean intensity of each membrane.

### Prediction of Lysine Ubiquitination

For lysine ubiquitination prediction we used the Rapid UBIquitination detection (RUBI) version 1 tool using 1 % and 5 % false positive rate by the Department of Biomedical Sciences, University of Padova^39^.

### Protein Sequence Analysis

To predict protein parameters including pI, amino acid composition and charge we used the Expasy ProtParam online tool operated by the SIB Swiss Institute of Bioinformatics^40^. Conservation level was determined using standard alignment parameters of the Jalview software (RRID:SCR_006459) developed in Geoff Barton’s Group, Division of Computational Biology, School of Live Sciences at the University of Dundee, Scotland, UK^41^.

### Protein Structure Prediction

AlphaFold models AF-Q86W26-F1-model_v6 and AF-Q8CCN1-F1-model_v6 were used and models were visualised using the UCSF ChimeraX software (RRID:SCR_015872) version 1.9 (2024-12-11)^42–44^.

### Data Visualisation and Analysis

Experimental data were visualised using GraphPad Prism 7 software and always represent Mean ± S.E.M. and statistics were performed using the integrated statistic tools. Vector graphics were generated with Inkscape (v1.4 (86a8ad7, 2024-10-11)).

## Acknowledgements

We thank Dr. Nora Mirza, Rebekka Bauer, Yvonne Postma, Lucy Biber (née Hezinger) and Sophia Zagar for help with the generation of plasmids, stable cell lines, and CRISPR/eCas9 knockout of NLRP10. We thank Susanne Reisse, head of the core imaging unit of the core facilities at the University of Hohenheim (CFH-IMG), for excellent supervision at the light microscope. This work was supported by a grant from the DFG to M.G. (GE 976/16-1). Parts of the equipment used were supported by the EFRE EU fund (grant no. 2172959). Publishing fees are supported by Funding Programme Open Access Publishing of University of Hohenheim. M.G. is supported by the European Research Council (ERC Advanced Grant NalpACT) and by the DFG under Germany’s Excellence Strategy–EXC2151-390873048.

## Conflict of Interest

All authors declare that they have no competing financial interests or personal relationships that could have appeared to influence the work reported in this paper.

## Notes

### Competing Interest Statement

The authors have declared no competing interest.

