## Supplementary Information for "C-terminal lysine residues localise NLRP10 at lipid compartments and govern NLRP10 oligomer formation"

[1]Department of Immunology, Institute of Nutritional Medicine, University of Hohenheim, Stuttgart, Germany
[2]Institute of Structural Biology, University of Bonn, Venusberg-Campus 1, Bonn, Germany
[3]present address: Institute of Nutritional Medicine, University Hospital Schleswig-Holstein, Campus Lübeck, University of Lübeck, Lübeck, Germany

The PDF file includes:

- Supplementary Figures S1 – S6
- Supplementary Tables S1 – S3

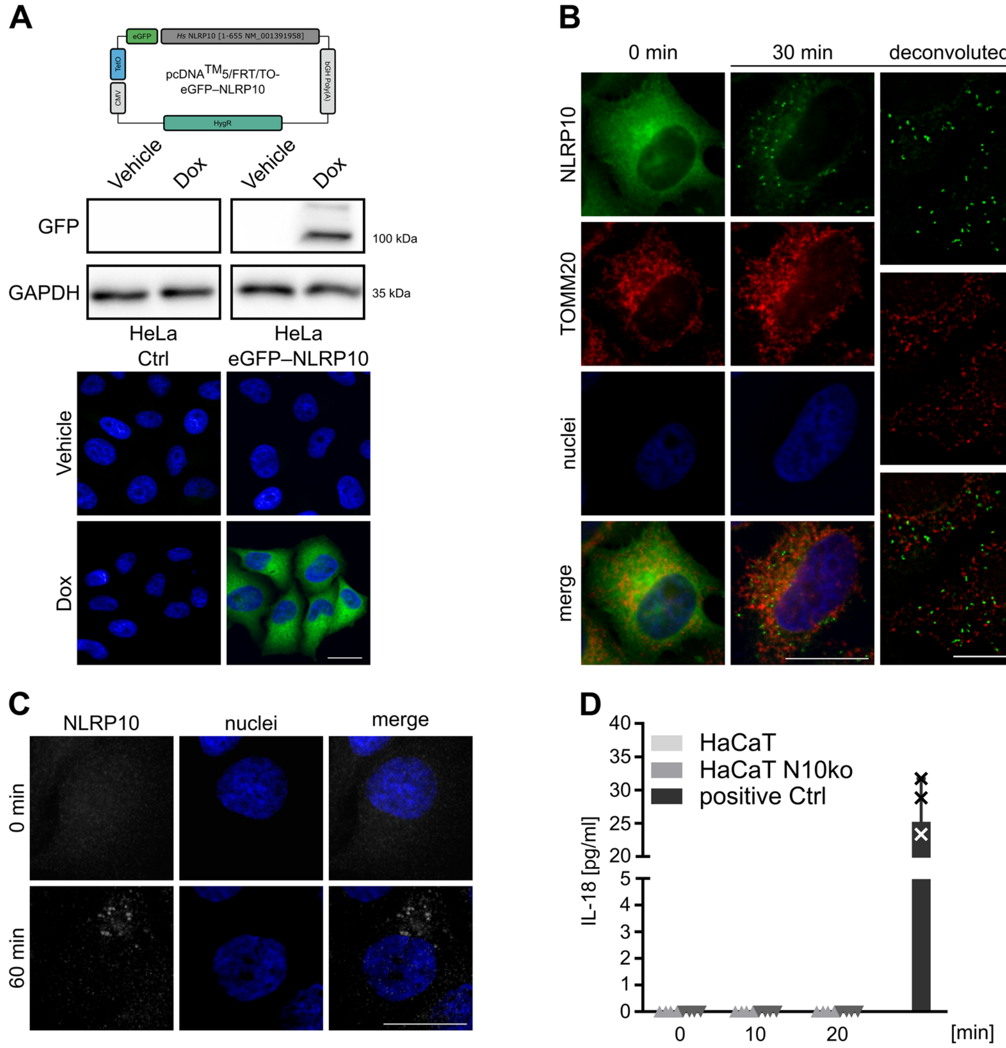


**Supplementary Fig. S1 | HeLa and HaCaT cells respond with *m*-3M3FBS-dependent NLRP10 oligomerisation without co-localisation with mitochondrial marker TOMM20.** (**A**) Stable HeLa cells show expression of eGFP-NLRP10 only in response to Dox. Middle panel shows immunoblot analysis with antibodies directed against GFP and GAPDH as loading control, lower panel shows micrographs of cells treated with or without Dox. (**B**) Micrographs of eGFP-NLRP10 (green) expressing HeLa cells treated with *m*-3M3FBS (85 µM) stained with an antoibody directed against TOMM20. Right panel shows deconvoluted stack. (**C**) Micrographs of HaCaT cells stained with an antibody directed against NLRP10 and treated with 85 µM *m*-3M3FBS. Nuclei were stained with HOECHST dye (**A-C**). (**D**) Quantification of the IL-18 release of HaCaT cells treated with *m*-3M3FBS (85 µM) or Poly I:C (10 µg/ml) as a positive control. Scale bar: 20 µm.

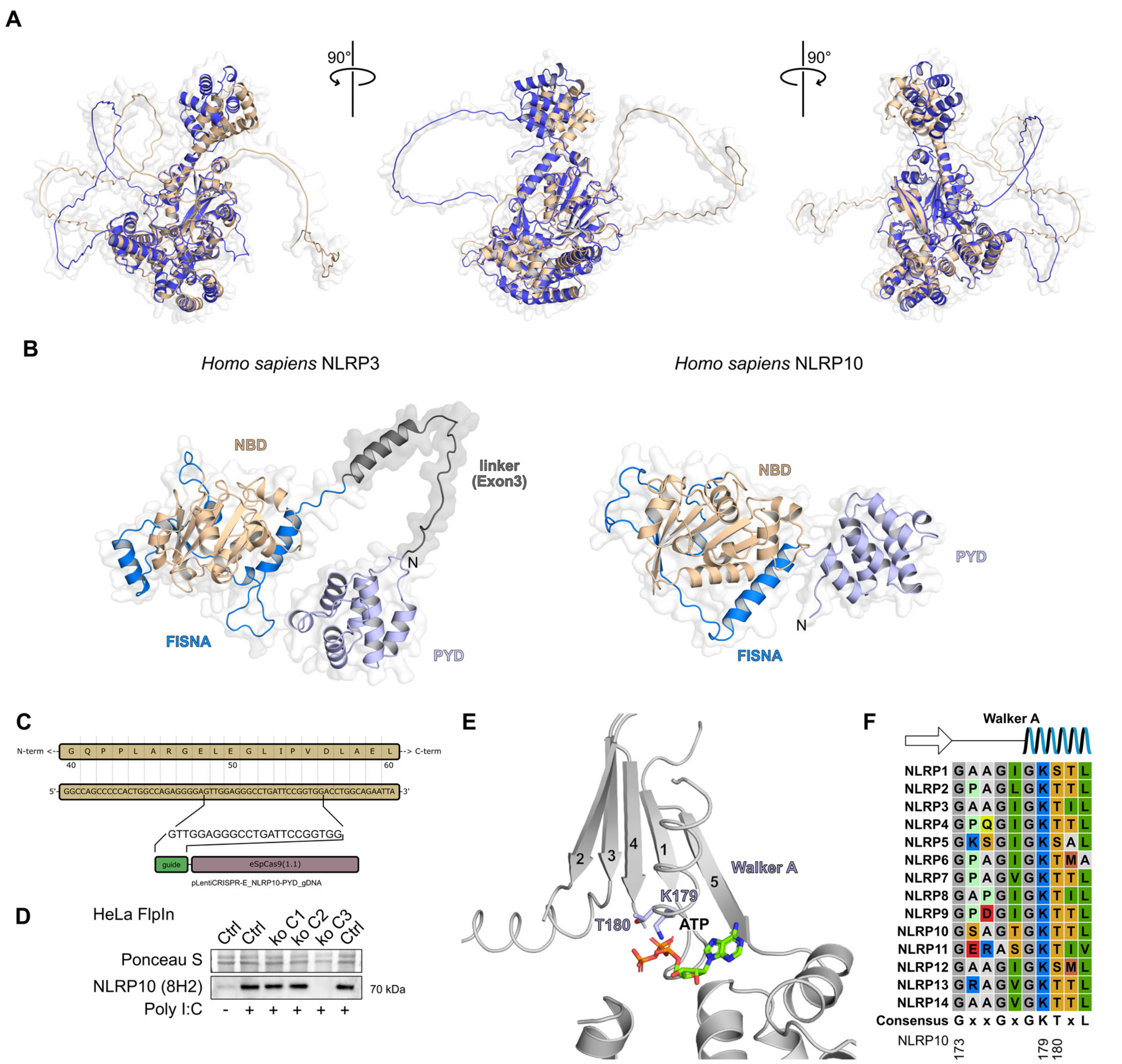


**Supplementary Fig. S2 | *Homo sapiens* and *Mus musculus* NLRP10 show high similarity with a different PYD-NACHT linker compared to NLRP3.** (**A**) Superpositioned AlphaFold models of human (blue) and mouse (beige) NLRP10 at three different angles. (**B**) The linker between the PYD and NACHT is encoded by an additional Exon in NLRP3, which results in a long stretch (grey), which is absent in NLRP10. (**C**) Schematics of the gDNA design to target the DNA anti-sense strand of the NLRP10 PYD. (**D**) Immunoblot analysis showing the established clone 3 of HeLa cells transfected with the pLentiCRISPR plasmid to target for NLRP10 knockout. 8H2 NLRP10 antibody has been used to detect NLRP10, whichs expression was enhanced using Poly I:C. (**E**) Cartoon of the nucleotide-binding site of NLRP10 modelled with AlphaFold3. The location of ATP refers to the model of NLRP3 (PDB: 7PZC). The Walker A motif is indicated with two catalytic active residues. (**F**) Sequence alignment of all NLRP family members with consensus for the Walker A motif.

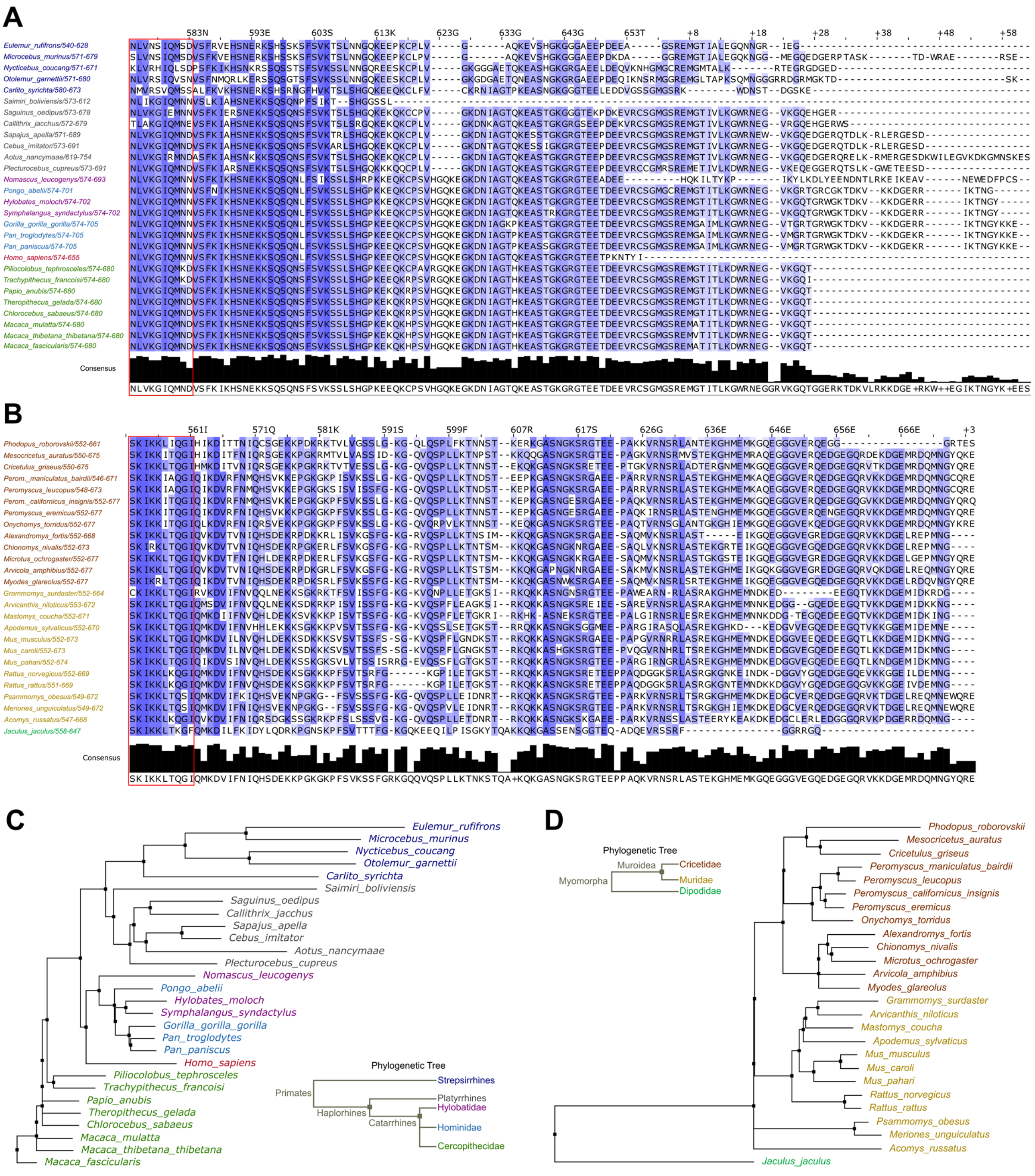


**Supplementary Fig. S3 | Evolution of the NLRP10 basic tail: *Homo sapiens* NLRP10 shows divergence among Haplorhines.** (**A-B**) Multi-sequence alignment of primate NLRP10 tails and myomorpha Nlrp10 tails. The last 10 amino acids of the HD2 are shown in red box. (**C-D**) Dendrogram according to NLRP10 sequence of primates or Nlrp10 sequences of Myomorpha. The dendrogram largely matches the general evolutionary distance of the species. The human NLRP10 (red) seperates from all other Catarrhines (**C**). Among Myomorpha, only *Jaculus jaculus* seperates, currently the only high quality predicted sequence for dipodidae (**D**). The colours of (**A-B**) match to (**C-D**) and Suppl. Tables **S1-S2**.

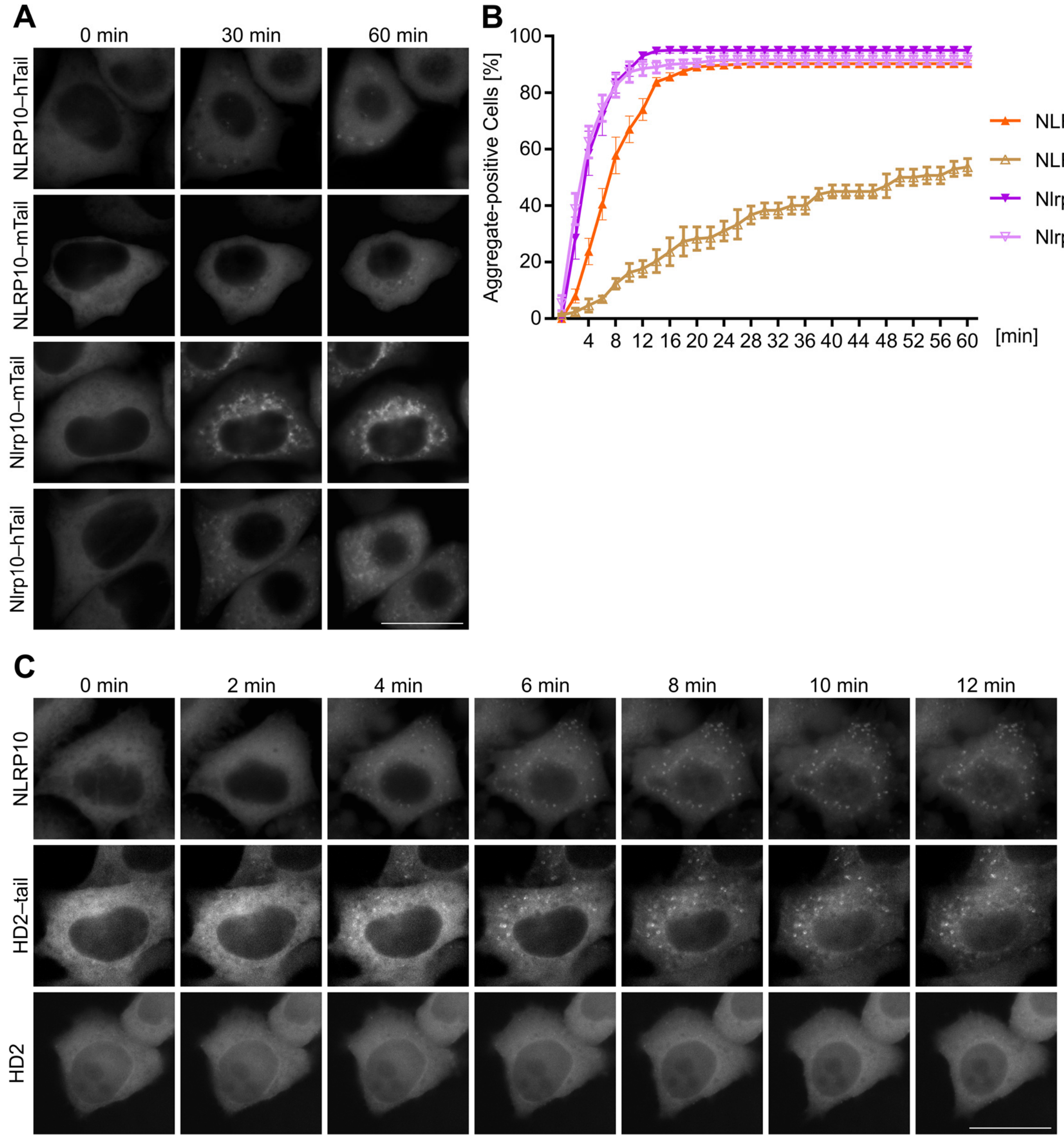


**Supplementary Fig. S4 | Human and mouse chimeric NLRP10 can aggregate and the tail is sufficient for is aggregation behaviour.** (**A**) Live-cell imaging micrographs of HeLa cells transiently transfected with chimeric NLRP10 and treated with *m*-3M3FBS (85 µM). (**B**) Quantification of the aggregate-positive cells from (**A**). (**C**) Live-cell imaging micrographs of HeLa cells treated with *m*-3M3FBS (85 µM), which were transiently transfected with eGFP fused to the HD2 and HD2–tail of NLRP10 compared to NLRP10. Scale bar 20 µm.

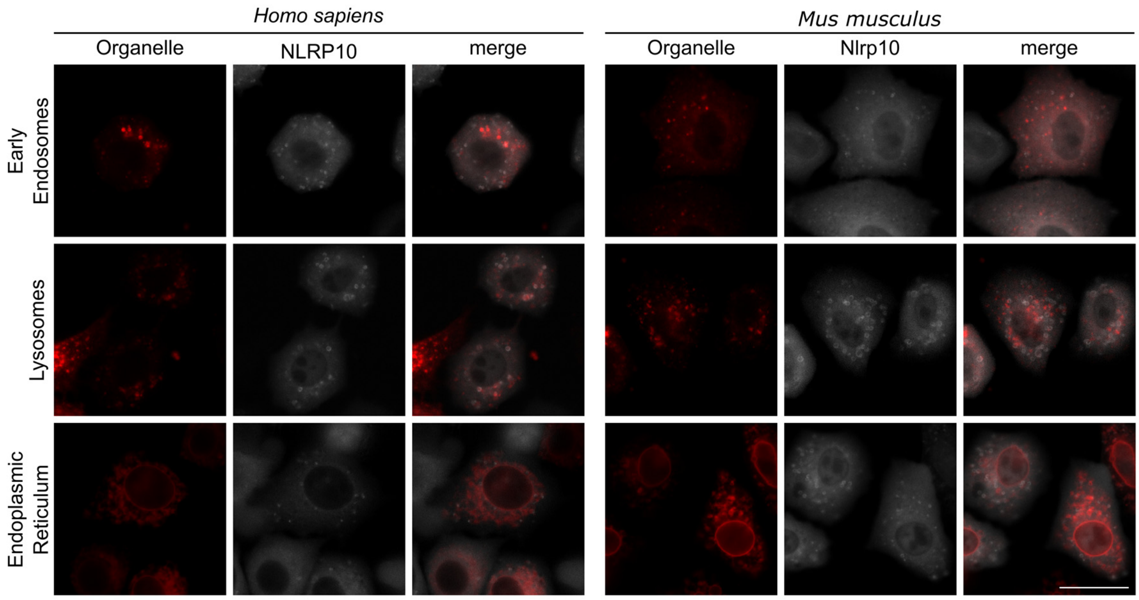


**Supplementary Fig. S5 | Human and mouse NLRP10 partially co-localised with early endosomes and lysosomes but not with the endoplasmic reticulum.** Live-cell imaging micrographs of HeLa cells expressing human NLRP10 (left) or mouse Nlrp10 (right) and treated with *m*-3M3FBS (85 µM). Scale bar 20 µm.

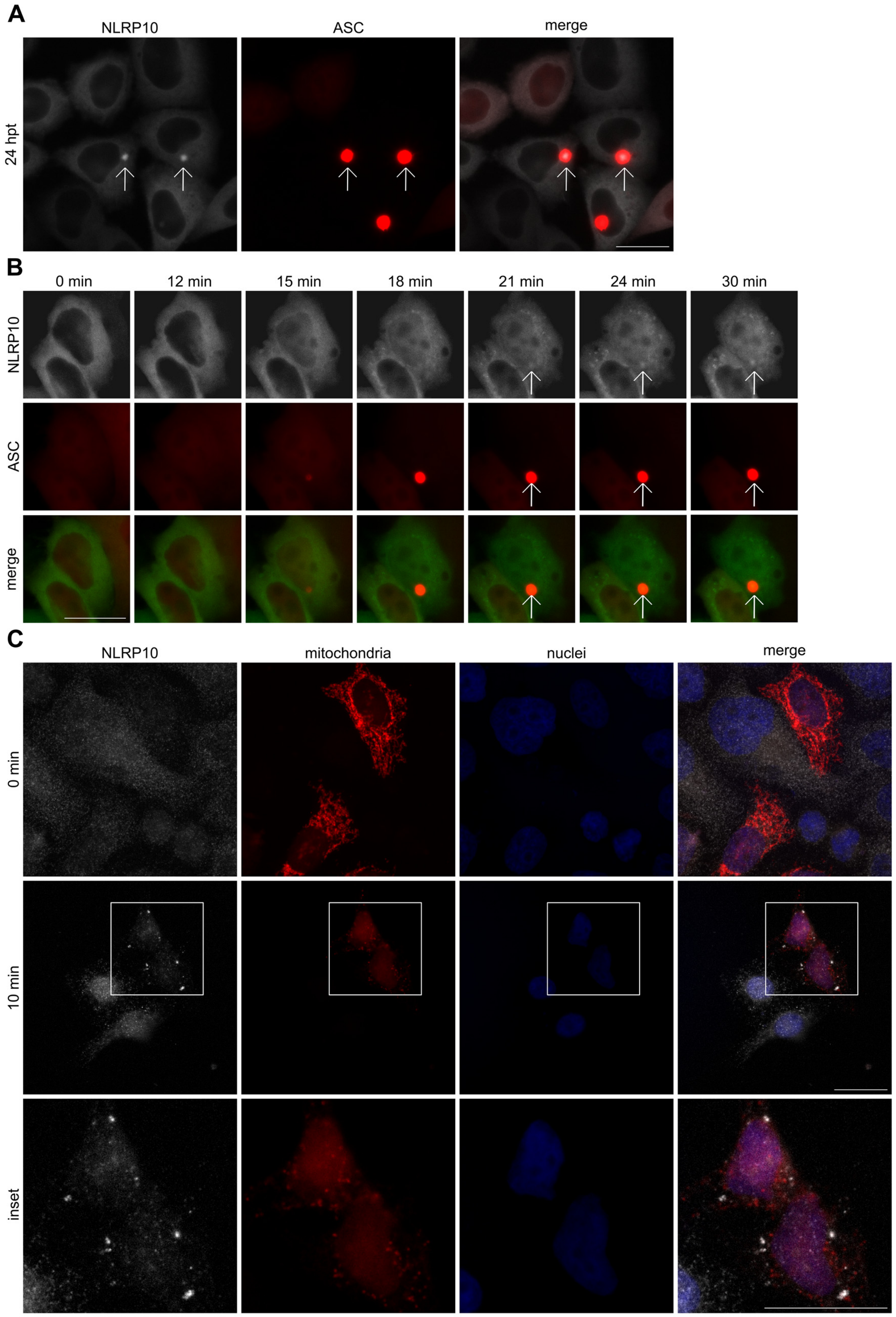


**Supplementary Fig. S6 | NLRP10 can form speck with ASC but ASC specks appear prior NLRP10 aggregation.** (**A**) Live-cell imaging micrographs of stable HeLa cells expressing eGFP-NLRP10 transiently transfected with ASC-RFP (24 h). (**B**) Live-cell imaging micrographs of stable HeLa cells expressing ASC-RFP under an CMVd1 promoter transiently transfected with eGFP-NLRP10 (24 h) and treated with *m*-3M3FBS (85 µM). (**C**) Micrographs of HeLa cells transiently transfected with mts-mCherry (red) and stained with an antibody directed againste NLRP10 (white) treated with or without *m*-3M3FBS (85 µM), nuclei were stained with HOECHST dye. White arrows indicate Specks. Scale bar 20 µm.

### Supplementary Tables

***Supplementary Table S1***: List of NLRP10 reference sequences of primates including total length and length of expected tail with expected pI according to human protein model and alignment mapping the last 10 amino acids of the *Homo sapiens* NLRP10 NACHT domain (NLVKGIQMNN).

| Reference | Species | Length | Tail (aa) | pI | Comment |
|---|---|---|---|---|---|
| XP_003920428.3 | *Saimiri boliviensis* | 612 | 30 | 10.00 | Shortest tail |
| NP_001378887.1 | *Homo sapiens* | 655 | 72 | 9.52 | |
| XP_069325622.1 | *Eulemur rufifrons* | 628 | 79 | 5.91 | Low tail pI |
| XP_008060833.2 | *Carlito syrichta* | 673 | 84 | 9.01 | Starts with T |
| XP_053415859.1 | *Nycticebus coucang* | 671 | 91 | 9.08 | |
| KAK2100778.1 | *Saguinus oedipus* | 678 | 96 | 9.55 | |
| XP_008005997.2 | *Chlorocebus sabaeus* | 680 | 97 | 9.25 | |
| XP_015290797.3 | *Macaca fascicularis* | 680 | 97 | 9.47 | |
| XP_001098361.3 | *Macaca mulatta* | 680 | 97 | 9.24 | |
| XP_050615200.1 | *Macaca thibetana thibetana* | 680 | 97 | 8.81 | |
| XP_021781824.2 | *Papio anubis* | 680 | 97 | 9.63 | |
| XP_023045669.1 | *Piliocolobus tephrosceles* | 680 | 97 | 8.52 | |
| XP_033061140.1 | *Trachypithecus francoisi* | 680 | 97 | 9.30 | |
| XP_025211582.1 | *Theropithecus gelada* | 680 | 97 | 8.93 | |
| XP_002755056.3 | *Callithrix jacchus* | 679 | 98 | 9.83 | |
| XP_012639395.1 | *Microcebus murinus* | 679 | 99 | 5.95 | Low tail pI |
| XP_003781555.1 | *Otolemur garnettii* | 680 | 100 | 9.66 | |
| XP_017355843.1 | *Cebus imitator* | 691 | 108 | 8.96 | |
| KAL0629402.1 | *Plecturocebus cupreus* | 691 | 109 | 7.86 | |
| XP_032096543.1 | *Sapajus apella* | 689 | 109 | 8.96 | |
| XP_030685543.1 | *Nomascus leucogenys* | 693 | 110 | 8.49 | |
| XP_012311817.2 | *Aotus nancymaae* | 754 | 117 | 9.39 | Atypical PYD |
| XP_002822118.2 | *Pongo abelii* | 701 | 118 | 9.76 | |
| XP_032021750.1 | *Hylobates moloch* | 702 | 119 | 9.84 | |
| XP_055140623.1 | *Symphalangus syndactylus* | 702 | 119 | 9.93 | |
| XP_055212467.2 | *Gorilla gorilla gorilla* | 705 | 122 | 9.81 | Longest tail |
| XP_003818292.1 | *Pan paniscus* | 705 | 122 | 9.73 | Longest tail |
| XP_003313013.5 | *Pan troglodytes* | 705 | 122 | 9.73 | Longest tail |

***Supplementary Table S2***: List of Nlrp10 reference sequences of myomorpha including total length and length of expected tail with expected pI according to mouse protein model and alignment mapping the last 10 amino acids of the *Mus musculus* Nlrp10 NACHT domain (SKIKKLTQGI).

| Reference | Species | Length | Tail (aa) | pI | Comment |
|---|---|---|---|---|---|
| XP_012805509.2 | *Jaculus jaculus* | 647 | 80 | 9.85 | Shortest tail |
| XP_051048548.1 | *Phodopus roborovskii* | 661 | 100 | 10.0 | |
| XP_028623966.1 | *Grammomys surdaster* | 664 | 103 | 9.88 | |
| XP_050019272.1 | *Alexandromys fortis* | 668 | 107 | 9.93 | |
| NP_001099761.1 | *Rattus norvegicus* | 669 | 108 | 9.43 | |
| XP_052011198.1 | *Apodemus sylvaticus* | 670 | 109 | 9.46 | |
| XP_032751552.1 | *Rattus rattus* | 669 | 109 | 9.55 | |
| XP_076796644.1 | *Arvicanthis niloticus* | 672 | 110 | 9.62 | |
| XP_031243649.1 | *Mastomys coucha* | 671 | 110 | 9.13 | |
| XP_051005138.1 | *Acomys russatus* | 668 | 112 | 9.79 | |
| XP_057635579.1 | *Chionomys nivalis* | 673 | 112 | 9.82 | |
| XP_021024703.1 | *Mus caroli* | 673 | 112 | 9.13 | |
| NP_780741.1 | *Mus musculus* | 673 | 112 | 8.82 | Lowest pI |
| XP_021074826.1 | *Mus pahari* | 674 | 113 | 9.22 | |
| XP_021517635.2 | *Meriones unguiculatus* | 672 | 114 | 9.52 | |
| XP_055463457.1 | *Psammomys obesus* | 672 | 114 | 9.63 | |
| XP_038199953.1 | *Arvicola amphibius* | 677 | 116 | 9.94 | |
| XP_035297287.1 | *Cricetulus griseus* | 675 | 116 | 9.43 | |
| XP_005075823.1 | *Mesocricetus auratus* | 675 | 116 | 9.43 | |
| XP_005351132.1 | *Microtus ochrogaster* | 677 | 116 | 9.73 | |
| XP_048316419.1 | *Myodes glareolus* | 677 | 116 | 9.77 | |
| XP_036033111.1 | *Onychomys torridus* | 677 | 116 | 9.99 | |
| XP_052619135.1 | *Peromyscus californicus insignis* | 677 | 116 | 9.65 | |
| XP_059107555.1 | *Peromyscus eremicus* | 677 | 116 | 9.88 | |
| XP_028750420.1 | *Peromyscus leucopus* | 673 | 116 | 9.76 | |
| XP_006978619.1 | *Peromyscus maniculatus bairdii* | 671 | 116 | 9.63 | |

***Supplementary Table S3***: List of oligonucleotide primers used in this study.

| Name | Sequence (5'->3') | Usage |
|---|---|---|
| NLRP10_F | CAAGGGATCCATGGCCATGGCC | N-term. |
| NLRP10_R | TAGACTCGAGTTATATGTAAGTATTTTTTGGTG | C-term. |
| NLRP10_1-583_R | GTGGTCGACGATATCTTAATTGTTCATCTGAATACC | Δ584 |
| NLRP10_1-482_R | GTGGTCGACGATATCTTACTCTTTCACCAGGTAAGACATGG | Δ483 |
| NLRP10_ΔPYD_F | GCGCGGATCCTACAGAGAAGTATACCGAGAGCATGTG | ΔPYD |
| Nlrp10_BglII_F | GAAGATCTATGGCCTTGGCACGGGCCAA | N-term. |
| Nlrp10_XhoI_R | GATCCTCGAGCTACCCATTCATC | C-term. |
| Nlrp10dPYD_BglII_F | GAAGATCTATGGAGCTTGTAGACTACCTCA | ΔPYD |
| Nlrp10Δtail_XhoI_R | GATCCTCGAGTTAGATACCTTGTGTGAGCTTCT | Δ562 |
| NLRP10_nsc_XhoI | TAGACTCGAGATTGTTCATCTGAATACC | chimeras |
| Nlrp10Δtail_nsc_XhoI_R | GATCCTCGAGGATACCTTGTGTGAGCTTCT | chimeras |
| XhoI_hTail_F | GATCCTCGAGGTATCATTCAAGATAAAACATTC | chimeras |
| hTail_ApaI_R | GTGGGCCCTTATATGTAAGTATTTTTTGGTG | chimeras |
| XhoI_mTail_R | GATCCTCGAGCAGATGAAAGATGTCATTCTC | chimeras |
| mTail_ApaI_R | CTGGGCCCCTACCCATTCATCTTATCTATCATC | chimeras |
